# Protein Expression Patterns in ovarian cancer cells Associated with Monofunctional Platinums Treatment

**DOI:** 10.1101/628958

**Authors:** Laila Arzuman, Mohammad Ali Moni, Philip Beale, Jun Q. Yu, Mark Molloy, Julian M.W. Quinn, Fazlul Huq

## Abstract

Platinum drugs cisplatin and carboplatin, given in combination with paclitaxel, constitute the standard chemotherapy against ovarian cancer (OC). Oc chemoresistance is a major obstacle to effective treatment, but knowledge of the mechanisms that underlie it remains incomplete. We thus sought to discover key proteins associated with platinum resistance by comparing A2780 OC cells with A2780^cisR^ cells (resistant cells derived from the A2780 line) to identify proteins with markedly altered expression levels in the resistant cells. We also determined which proteins in these cells had altered expression in response to treatment with either designed monofunctional platinum alone or a combination with cisplatin with selected phytochemical therapeutic agents.

We thus performed proteomic analysis using 2D-gel electrophoresis A2780 and A2780^cisR^ to identify proteins with differential expression; these were eluted and analysed by mass spectrometry to identify them. A total of 122 proteins were found to be differentially expressed between A2780 and A2780^cisR^ cell lines in the absence of any drug treatment. Among them, levels of 27 proteins in A2780^cisR^ cell line were further altered (up-or down-regulated) in response to one or more of the drug treatments. We then investigated primary OC tissue RNA expression levels (compared to l ovarian tissue) of genes coding for these candidate 27 proteins using publically available datasets (The Cancer Genome Atlas). We assessed how expression of these genes in OC tissue associates with patient survival using Cox Proportional Hazard (PH) regression models to determine relative risk of death associated with each factor. Our Cox PH regression-based machine learning method confirmed a significant relationship of mortality with altered expression of ARHGDIA, CCT6A and HISTIH4F genes. This indicated that these genes affect OC patient survival, i.e., provided mechanistic evidence, in addition to that of the clinical traits, that these genes may be critical mediators of the processes that underlie OC progression and mortality.

Thus, we identified differentially expressed proteins that are implicated in platinum-based chemotherapy resistance mechanisms which may serve as resistance biomarkers. These drug resistance associated proteins may also serve as potential OC therapeutic targets whose blockade may enhance the effectiveness of platinum based drugs.

## 1 BACKGROUND

Ovarian cancer (OC) is the leading cause of death from gynaecological cancers in women [15, 36, 29] and the fifth most common cause of cancer deaths in women overall, with an estimated 239,000 new cases and over 152,000 deaths worldwide annually [10, 27]. It has a five year mortality rate in the United States of approximately 35% [12, 35] [3, 22]. Current standard treatment mainly comprises platinum-based chemotherapeutics, with platinum drugs cisplatin and carboplatin given in combination with paclitaxel used to treat nearly all women with OC. However, the most prevalent OC type, high grade serous carcinoma (HGSC), often develops significant chemotherapy resistance alongside greatly altered genomes and transcriptome. Thus, most patients initially respond to the treatment, but over the longer term few are cured [13, 17].

We have previously designed a number of novel platinum and palladium complexes with the aim of overcoming platinum resistance in ovarian cancer [ref Huq et al], demonstrating that these show significantly greater activity than cisplatin in resistant tumour models [28]. We also studied a number of combinations of platinum drugs used in the clinic in complexes with tumour active phytochemicals, finding that a number of the combinations show cooperatively increased activity in ovarian tumour models. However, to systematically address the problem of cisplatin resistance we need to understand the alterations in cellular pathways that underlie this phenomenon, and endow cells with a survival advantage in patients being treatment. Such pathways may enable the cell to ignore growth inhibitory signals, to inappropriately block pro-apoptotic signals or to survive toxic chemical effects of the drug until serum levels of the drug fall [21, 25].

One-third of women treated with cisplatin develops resistance to the drug, and almost all of ovarian cancer (OC) patients need treatment for recurrent disease. Identification of biomarkers is essential for the resistant tumours towards the development of better therapies to improve the outcomes. Differential expression of proteins in platinum-resistant and -sensitive OC cells are believed to be significantly involved in pathways that are responsible for modulation of cisplatin action and hence are functional candidates for both therapeutic targets and treatment response biomarkers.

We therefore sought protein expression that was altered in cisplatin-resistant cells. To achieve this we undertook proteomic analysis of two closely related OC cell lines, A2780 and A2780^cisR^. Proteins thus identified were A2780 and A2780^cisR^ cells were then studied to determine responses to treatment with designed monofunctional platinum alone [6, 5, 23] and with out previously characterised combination of cisplatin and phytochemicals. We thus identified chemotherapy resistance-associated pathways and subsequent analysis of key pathway proteins using clinical transcriptomic data from OC patients showed that transcripts of some of these proteins are associated with patient surviva. We determined this using the rich datasets available from The Cancer Genome Atlas (TCGA) project, a collaboration between the National Cancer Institute (NCI) and National Human Genome Research Institute (NHGRI). The influence of clinical factors and disease marker gene expression on OC patient survival were determined using standard Cox Proportional Hazard (CPH) ([9]) models. Our analysis followed the methodology of Xu and Moni (2015) ([40, 26]) who used Cox PH regression modelling for the analysis of post-diagnosis survival time.

## 2 MATERIALS AND METHODS

### 2.1 Chemicals

Cisplatin (Cis) was prepared following Dhara’s method [4, 31]. LH3 and LH4 were synthesized according to previously described methods [6, 5]. Stock solutions of LH3, LH4, curcumin, genistein, capsaicin and quercetin were prepared as described previously [6, 5]. Briefly, solutions of LH3 and LH4 were made in 2:1 mixtures of DMSO and mQ water. Phytochemicals curcumin, genistein and quercetin were dissolved in ethanol to attain 1mM concentration whereas quercetin was dissolved in 1: 4 mixtures of DMF and mQwater.

### 2.2 Cell lines

Two ovarian cancer cell lines A2780 and A2780^cisR^ used in this study were provided by Dr Philip Beale (NSW Cancer Centre, Royal Prince Alfred Hospital, Sydney, Australia [14, 7, 24]. At first, A2780 (the parent cell line) was established from non-treated fresh tissue of ovarian cancer patient from which cisplatin resistant A2780^cisR^ cell line was obtained by the gradual exposure of A2780 cells with increasing doses of cisplatin until A2780 became resistant to the drug [11, 33, 34].

### 2.3 Proteomic study of the OC cell lines

Proteomics were performed to determine the proteins that underwent changes in expression in A2780 and A2780cisR cell lines before and after treatment with LH3 alone and its synergistic combinations with Cur (0/0 h) and Gen (0/0 h), and with LH4 alone and its synergistic combinations with Caps (4/0 h) and Quer (0/0 h) for 24 h at the median effect dose. Cell pellets were collected before and after drug treatments, washed with ice cold PBS, centrifuged for 2 mins at 3500 rpm at 4°C. The solution used for cell lysis consisted of 8M Urea, 4% CHAPS, 2 M thiourea and 65 mM dithiothreitol (BIORAD, Australia). Isoelectric focusing (IEF) for samples was performed using pH 3-10 non-linear ReadyStripTM IPG Strip, 11 cm, in Protean i12 IEF cell unit (BIORAD, Australia) for which samples were rehydrated in 60 mM dithiothreitol. 0.2% carrier ampholyte, 0.0002% bromophenol blu, 2 Mthiourea Murea, 4% CHAPS and deStreak (BIORAD, Australia). Two equilibration steps were carried out in SDS equilibration buffer consisting of SDS, 50% glycerol, 6 Murea, 1.5 M Tris HCl (pH 8.8), and bromophenol blue and 0.5 g Dithiothretiol in the first and 0.5 g iodoacetamide in the second. The concentration of protein was determined on Bio-Rad Protein Assay (BIO-RAD, Australia). SDS-PAGE was carried out by using 4-20% SDS CriterionTM TGXTM pre-cast gels in a cell separation unit Criterion DodecaTM (BIO-RAD, Australia), in the buffer containing Tris-glycine-HCl at 200 V for 100 min. Gels were stained in duplicate for 60 min with Bio-Safe Coomassie Stain (BIO-RAD, Australia). Melanie version 7.0 software (GeneBio, Swizterland) was used to analyse gel images obtained by ChemiDocTM MP Imaging System (BIO-RAD, Australia). A protein spot was considered to have undergone a significant change in expression if the fold change was greater than 1.5. Protein spots from preparative 2-D gels stained with Bio-Safe comma stain were washed with 120 l solution (50% acetonitrile (ACN)/50 mM NH4HCO3) and slowly agitated for half an hour at 37°C. The gels were washed with 25 I CAN and dried for 15 mins. The solution was removed, then spots were dried at 37°C for 15 mins followed by cooling at 4°C. The spots were digested for 10 min with 10 l trypsin held on ice[**?**].

Supernatants were transferred onto 96-well plate at 4°C to which was added 10 l of 25 mM NH4HCO3 for overnight digestion at 37°C. 0.1% trifluoroacetic acid (TFA) was used to extract peptides and concentrated by C18 zip-tips (Millipore,-C18, P10 size) on Xcise (Proteome Systems). A 1 ul aliquot was manually spotted on a MALDI AnchorChip plate with 1 l of a matrix made from 1 mg/mL in 90% v/v ACN, CHCA and 0.1% TFA. It was then air dried. MALDI-MS (Matrix-assisted laser desorption ionisation mass spectrometry) was done with 4800 plus MALDI TOF/TOF Analyser (AB Sciex). Samples were irradiated with a Nd:YAG (neodymium-doped yttrium aluminum garnet) and MS range of 700-4000 Da was used to obtain spectra. The eight strongest peptides from MS scan were isolated and fragmented by CID (collision induced dissociation) while instrument mode was set to MS/MS (TOF-TOF). The masses and intensities were measured following re-acceleration. Mascot (Matrix Science Ltd, London, UK) database search program was used to analysis the peptides masses data. SwissPort database was used to search the peak lists against Homo sapiens entries [16].

### 2.4 Survival association of transcripts of the identified proteins

We collected the RNAseq data for this study from TCGA genome data analysis centre (http://gdac.broadinstitute.org/) an interactive data system containing cancer genomic data [39]. We retrieved anonymized clinical data and RNAseq data for OC (Ovarian Serous Cystadeno Carcinoma TCGA, Provisional) from the cBioPortal ([8]).

In Clinical dataset there are 577 cases with 87 features. Cases that had RNAseq gene expression data included 535 cases with 19760 genes. We employed genes corresponding to the identified 27 significant proteins from the proteomics study.

Using these data we investigated a single outcome variable, namely OC-specific survival. 48 OC patient records that did not include clinical information were excluded from this analysis. We matched patient ID in both clinical and RNAseq dataset and identified 529 patients with data available for both. Among the clinical variables six clinical variables given above were considered ([20, 19]). Tumour histological subtype and age at initial diagnosis were collected from pathology reporting. OC stage was recorded according to American Joint Committee on Cancer (AJCC) staging classification ([1]). Patient survival data was taken from overall number of months of patient survival. Normal and tumour samples were identified using the TCGA barcode; two digits at position 14-15 of the barcode denote the sample type.

Tumour type spans from 01 to 09, normal type from 10 to 19 and control samples from 20 to 29. For gene expression, this was expressed and Fold change (FC) and variation in read count for individual genes accross tested individuals ([2]). FC is most suitable when the gene expression distribution is symmetric. However, in RNAseq analyses, expression levels are modelled by the discrete counts of reads mapped to a known gene in a reference genome. Poisson and Negative binomial distribution assumptions are taken for reading counts. When the abundance rate of a particular gene expression level is very low, read count distributions modelled by the Poisson or Negative binomial are skewed to the right. Thus, as using FC as a measure of differential expression may not be appropriate in such cases we transformed the gene expression value using a standardizing transformation and calculated z-scores for each expression value. For the expression from RNAseq experiments, the standard rule is to compute the relative expression of an individual gene in tumour samples using gene expression distribution in a reference population. A reference population was considered either all diploid tumours for the gene in question or, when available, normal adjacent tissue. The resulted value (z-score) indicates the number of standard deviations away from the mean expression in the reference population. Here, we considered genes showing *z* > ±1.96 to be differentially expressed. We computed z-scores for RNAseq data using following formula:

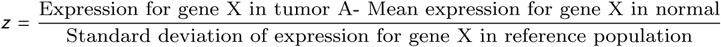

We thus used the z-score values to define samples with “altered” and “normal” (unaltered) expression of a gene. We have assumed a sample to be altered if the z-score for that sample is equal to or higher than a specific threshold value such as z=2, as noted. We therefore define altered versus normal as follows:

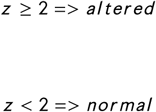

We performed the following analysis related to survival of patients with OC. First we used product-limit estimator for estimation of survival function, then we performed log rank test to determine whether the survival functions of two different groups (patients with altered gene expression and patients with unaltered gene expression) exhibit statistically significant difference and after then we used Cox PH regression models to determine the significance of genes, clinical factors on comparative risk of death and finally we performed functional analyses of our most significant genes found from our analysis.

We selected important clinical and demographic variables affecting OC of the corresponding genes of the 27 proteins that having experimentally determined significance in the proteomics study. The expression z-score for each gene was identified as being in “altered” or “normal” categories based on a significance threshold value (*z* < 2) as noted in the Data section above. We performed Cox PH regression for every gene individually known as univariate regression.

Survival analysis is a statistical analysis for estimating expected duration of time until one or more events happen, such as death in cancer and failure in mechanical systems. Normally, survival analysis is carried in three steps: determining time to event, formulating a censoring indicator for subject inclusion in the analysis, and the time to occurrence of event. Censoring in survival analysis is usually done in two ways, right censoring and left censoring. Right censoring occurs when a subject leaves the study before an event occurs, or the study ends before the event has occurred. Left censoring happens when the event of interest has already occurred before enrollment. This is very rarely encountered. Right censoring is again of two types. First one is Type I right censoring results from completely random dropout (e.g., emigration) and/or end of study with no event occurrence and the second one, Type II right censoring, occurs with end of study after fixed number of events amongst the subjects has occurred.

In survival analysis, one is interested in estimation of survival function of samples divided into subgroups or as whole. There are two types of estimation, one being non parametric in which no prior assumption of distribution of survival time is made and other being parametric estimation in which there are some assumption along with assuming predefined distribution of survival time. As noted above, we used non parametric technique for estimation of survival function. There are several such techniques, we used product-limit estimator. In short product-limit (PL) estimator of the survival function is defined as follows:

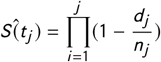

Here *Ŝ* (*t*_*j*_) is estimated survival function at time *t*_*j*_, *d*_*j*_ is the number of events occurred at *t*_*j*_, and *n*_*j*_ is the number of subjects available at *t*_*j*_. After estimating survival function, two or more groups can be compared using log-rank test. For example, we used Log-rank test to detect the most significant genes in the case of patient’s survival time in altered versus unaltered groups in context of gene expression. The null hypothesis is following: *H*_0_: Survival function for patients with altered gene expression is not different from the patients with normal (unaltered) gene expression

*H*_*A*_: Survival functions are different for these two groups. Symbolically these can be written

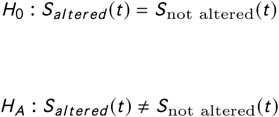

Using Cox PH regression, first we performed univariate survival analysis by selecting each gene separately, modelling the hazards of having the event under investigation (Ovarian Serous Cystadeno carcinoma, in this case), using an undetermined baseline hazard function and an exponential form of a set of covariates. Mathematically we can write the model as following:

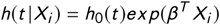

Whereas *h*(*t* |*X*) is the hazard function conditioned on a subject i with covariate information given as the vector *x*_*i*_, *h*_0_(*t*) is the baseline hazard function which is independent of covariate information, and represents vector of regression coefficients to the covariates correspondingly. We have calculated the hazard ratio (HR) based on the estimated regression coefficients from the fitted Cox PH model to determine whether a specific covariate affects patient survival. The hazard ratio for a covariate *x*_*r*_ can be expressed by the following simple formula exp (*β*_*r*_). Thus, the hazard ratio for any covariate can be calculated by applying an exponential function to the corresponding (*β*_*r*_) coefficient.

Pathways and gene ontology (GO) for these genes were analysed using KEGG pathway database (*http://www.genome.jp/kegg/p* and enriched using (*http://amp.pharm.mssm.edu/Enrichr/enrich*), a web based software tool.

## 3 RESULTS AND DISCUSSION

In this project, proteomics involving 2D-gel electrophoresis and mass spectrometry were carried out to identify the proteins that are differentially expressed in cisplatin-resistant A2780^cisR^ cell lines as compared to the levels found in cisplatin-sensitive A2780 cell line [37] and also to identify the proteins in the same cell line that underwent further changes in expression due to treatment with LH3 and its combinations with Cur and Gen administered as a bolus and with Caps and Quer administered using 4/0 h and 0/0 h sequences of administration. It was reported previously [6, 5] that the said combinations produced synergistic outcomes in the cell kill. 2D-gel images applying to A2780 and A2780^cisR^ cells before and after treatments are given in figure 1. A total of 122 spots were differentially expressed in A2780 and A2780^cisR^ cell lines before they were subjected to any drug treatment. Among them 27 spots in A2780^cisR^ cell line were significantly altered in expression (up-or down-regulated) due to treatments with LH3 and its combinations: LH3+Cur (0/0 h), LH3+Gen (0/0 h), as wells selected combinations of LH4: LH4+Caps (4/0 h) and LH4+Quer (0/0 h). Based on automatic spot detection by the software, a total of 122 spots were identified (Supplementary table 1). As mentioned earlier, the matched groups were finally grouped together into “Combine All” class for analysis on the differential protein expressions. From the same protein spots from different 2D-gel profiles that underwent changes in expression, nineteen matched groups were assigned by the Melanie software with the common matched ID numbers, as shown in Figures 1 and 2. Figures 3 to 7 show the protein profile applying to the drug treated A2780^cisR^ cell line.

**FIGURE 1.**
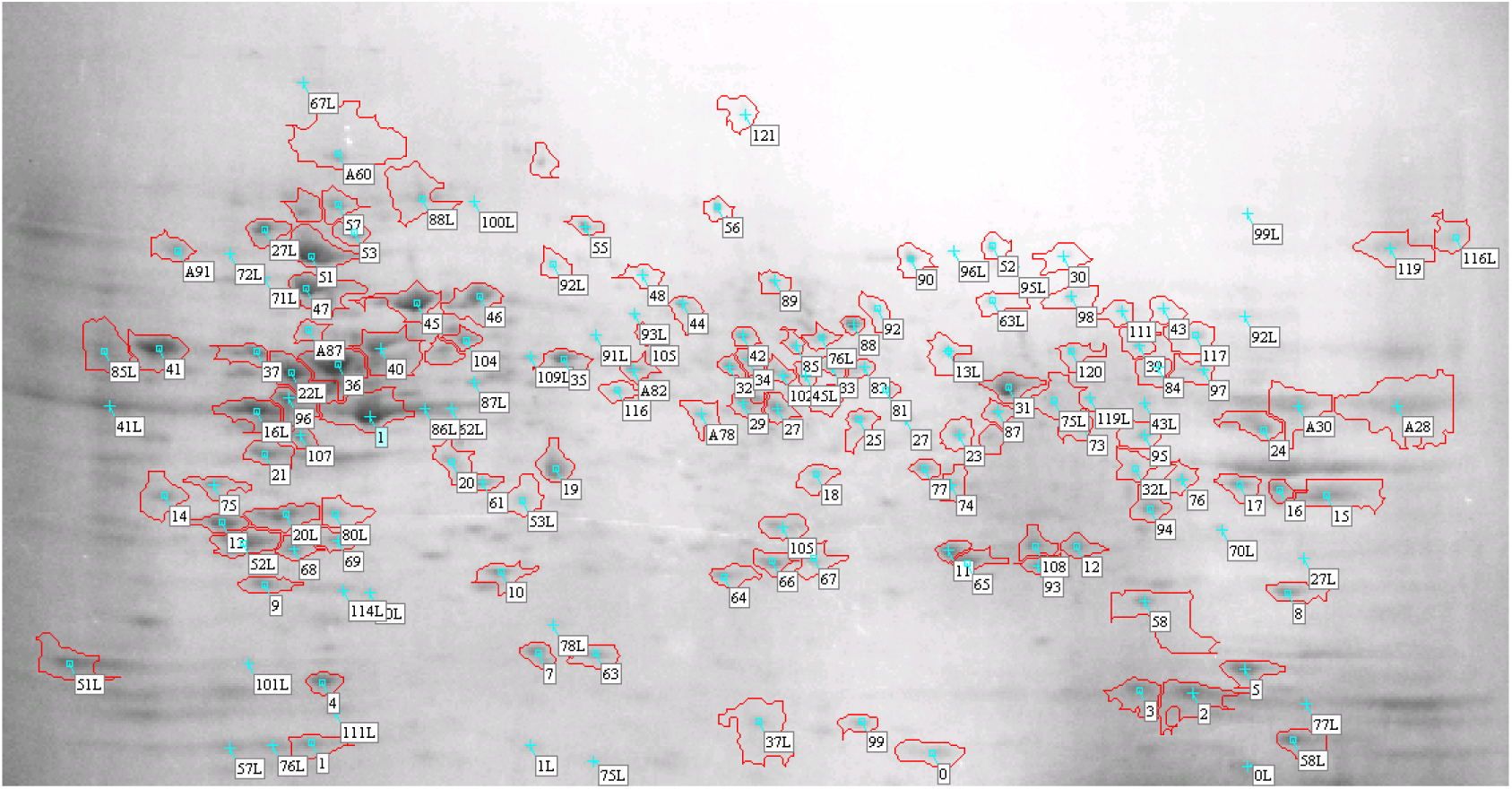
Protein profile of untreated A2780 cell line with annotated protein spots (along with Cut ID)

**FIGURE 2.**
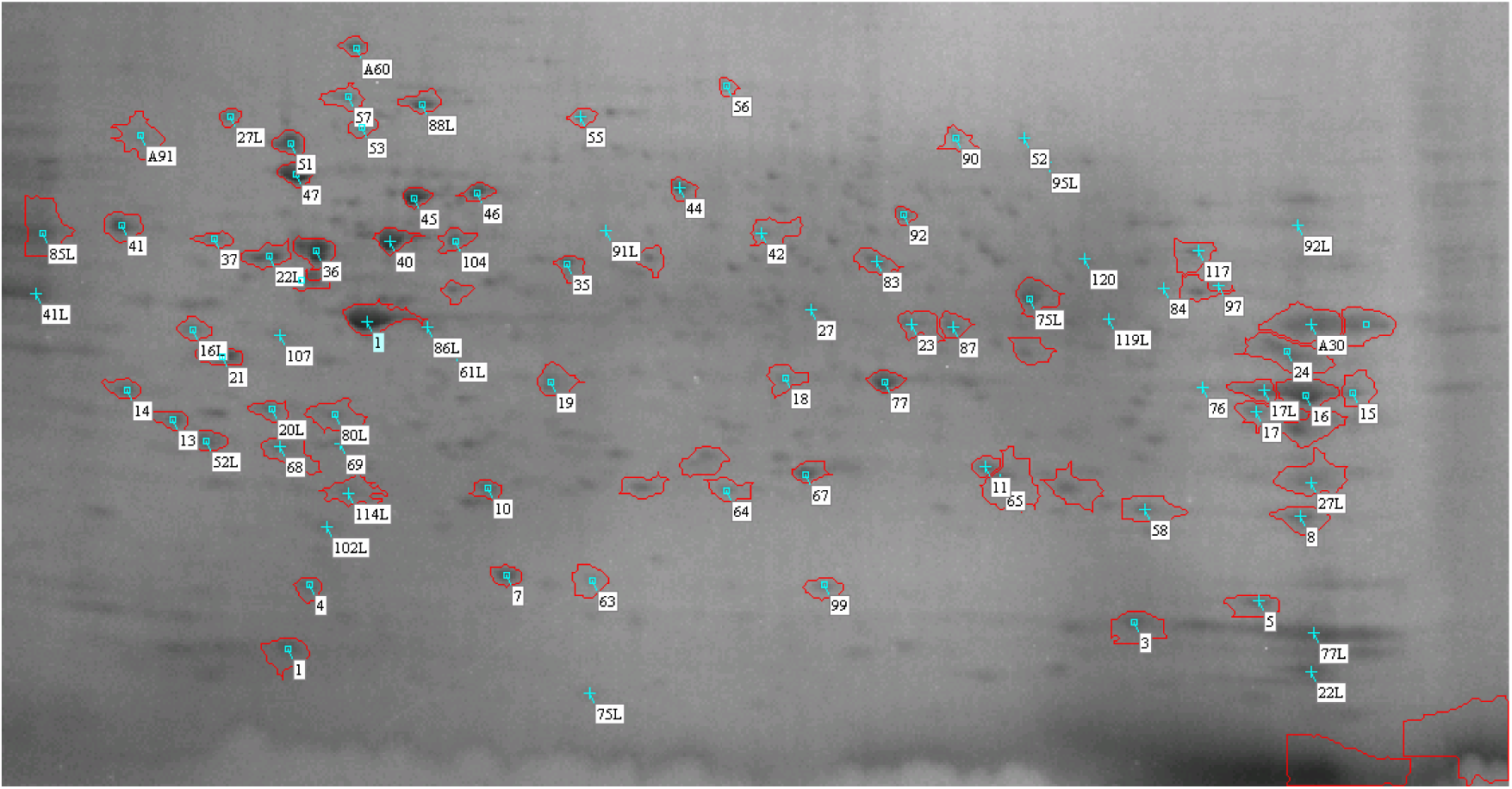
Protein profile of untreated A2780^cisR^ cell line with annotated protein spots (along with Cut ID) which have undergone changes in expression due to the selected drug treatments

**FIGURE 3.**
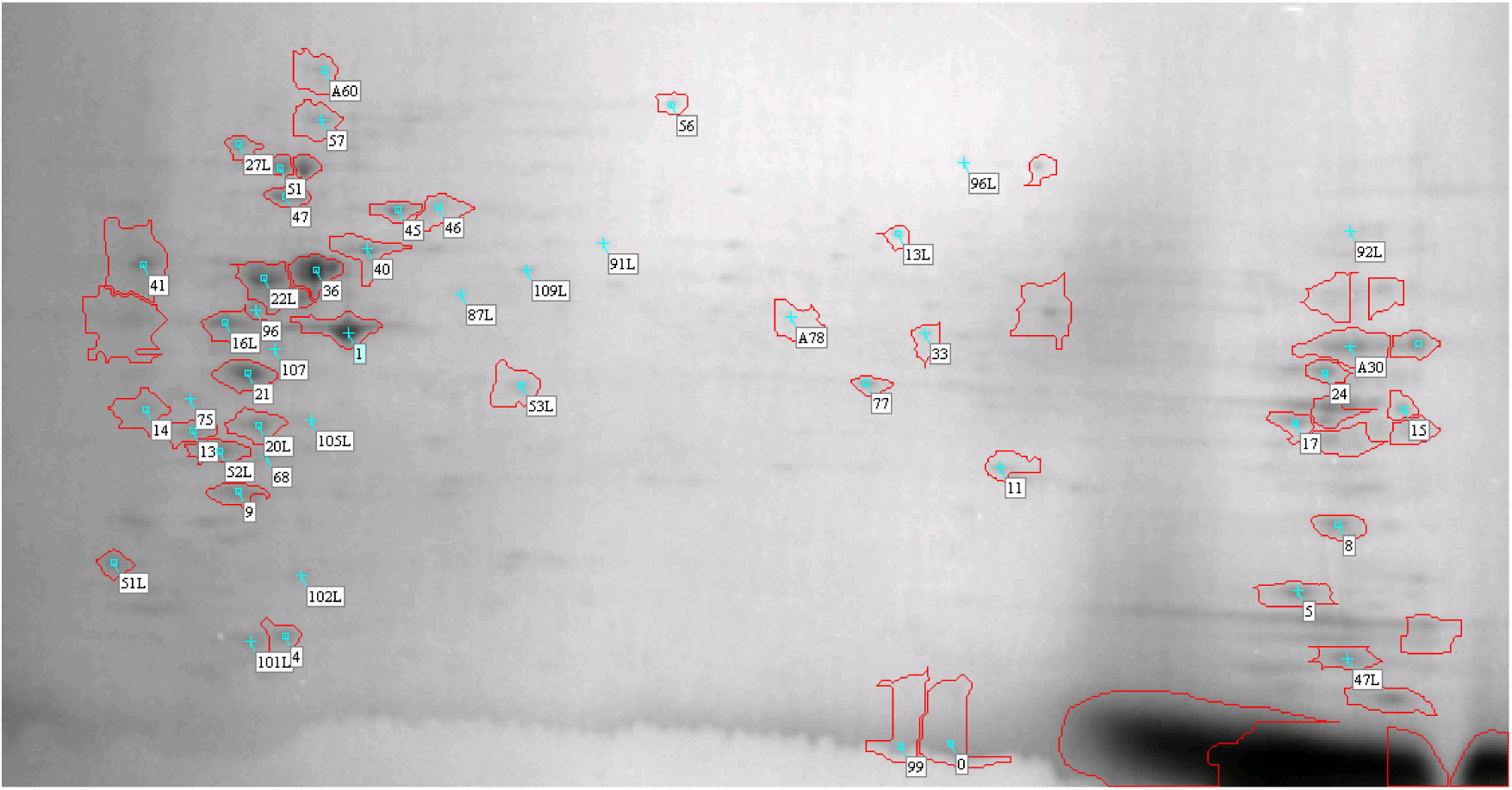
Protein profile of LH3 treated A2780^cisR^ cell line

**FIGURE 4.**
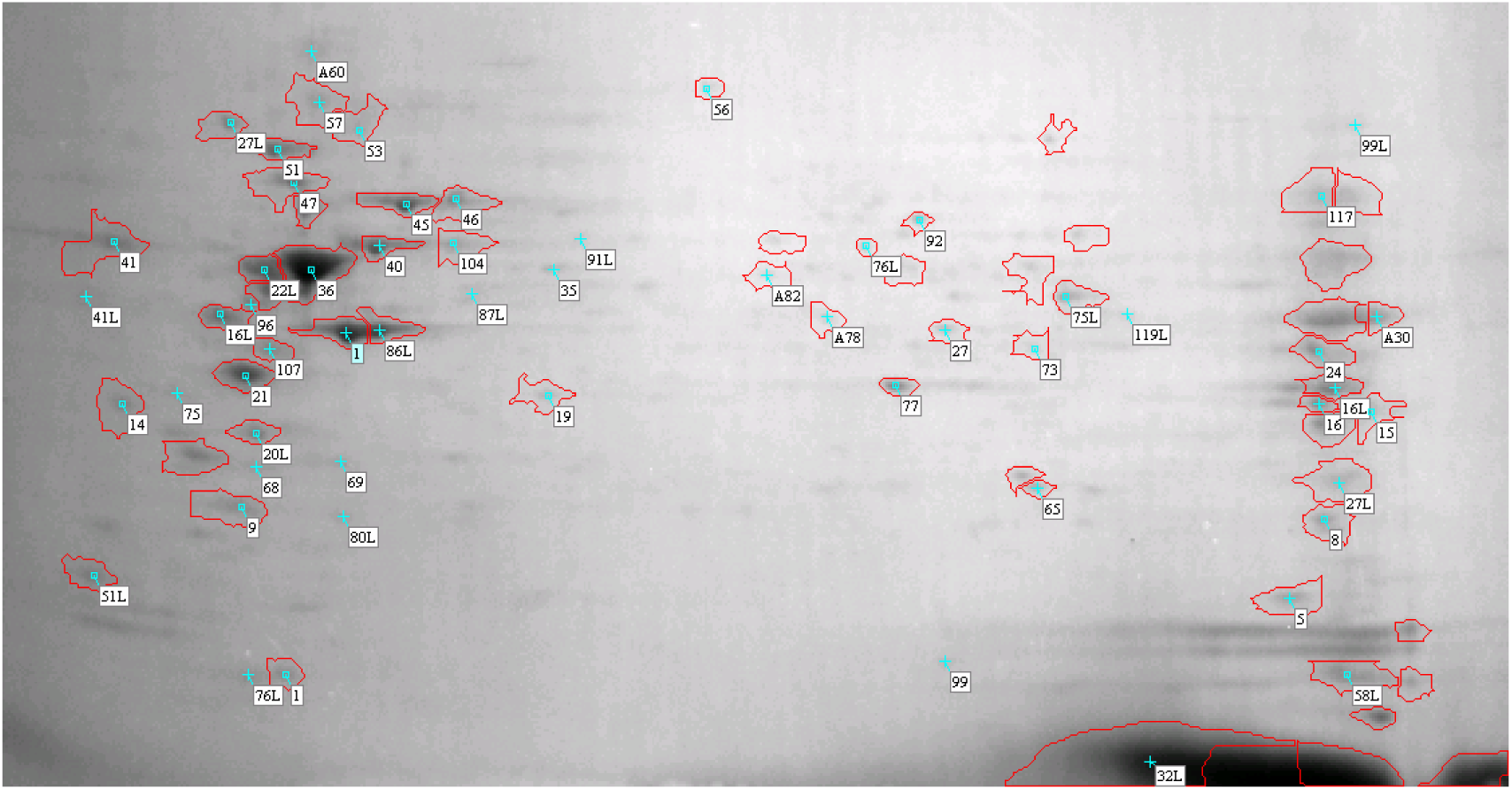
Protein profile of LH3+Cur (0,0) treated with A2780^cisR^ *cell line*

**FIGURE 5.**
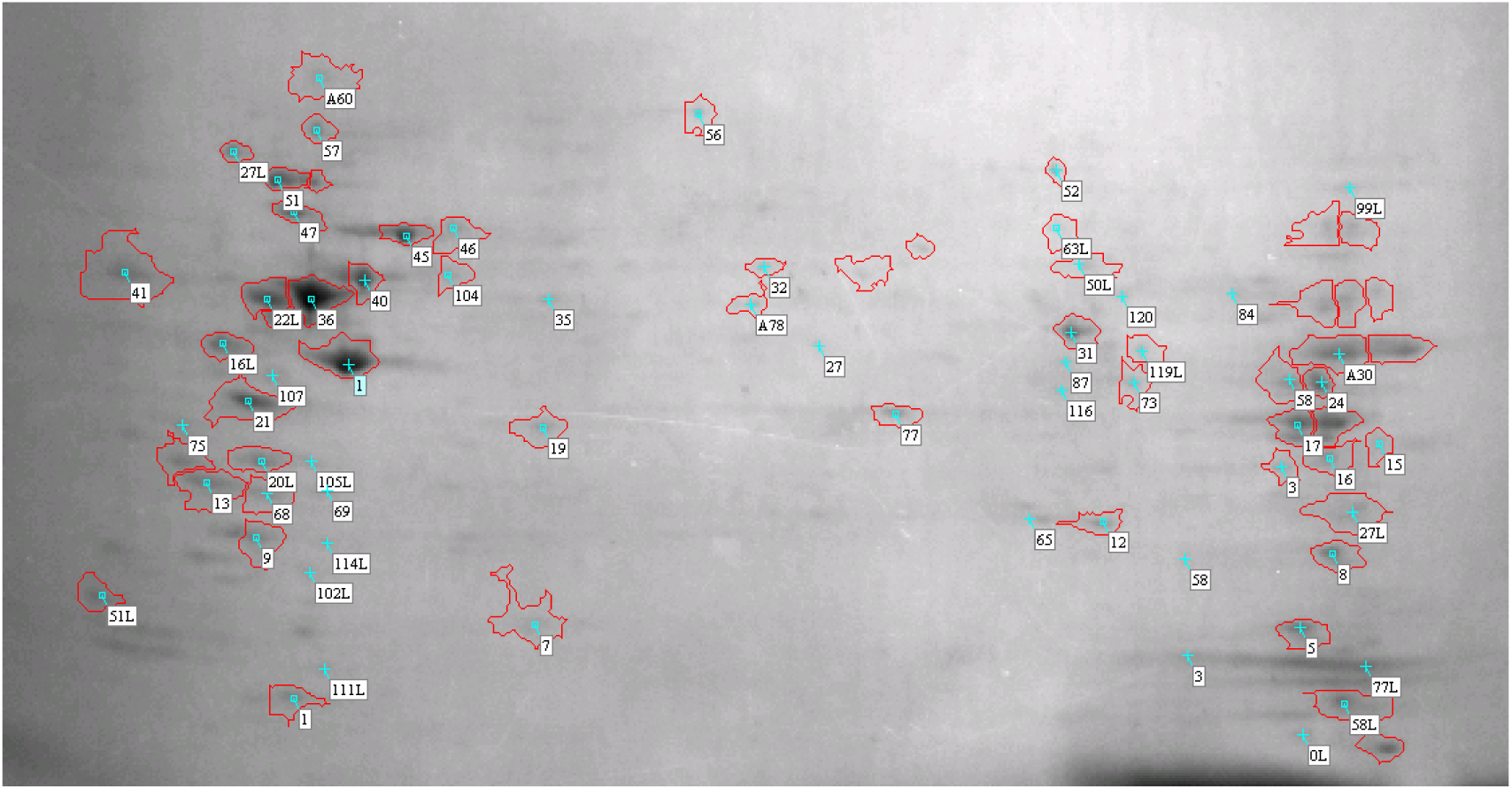
Protein profile of LH3+ Gen (0,0) treated with A2780^cisR^ cell line

**FIGURE 6.**
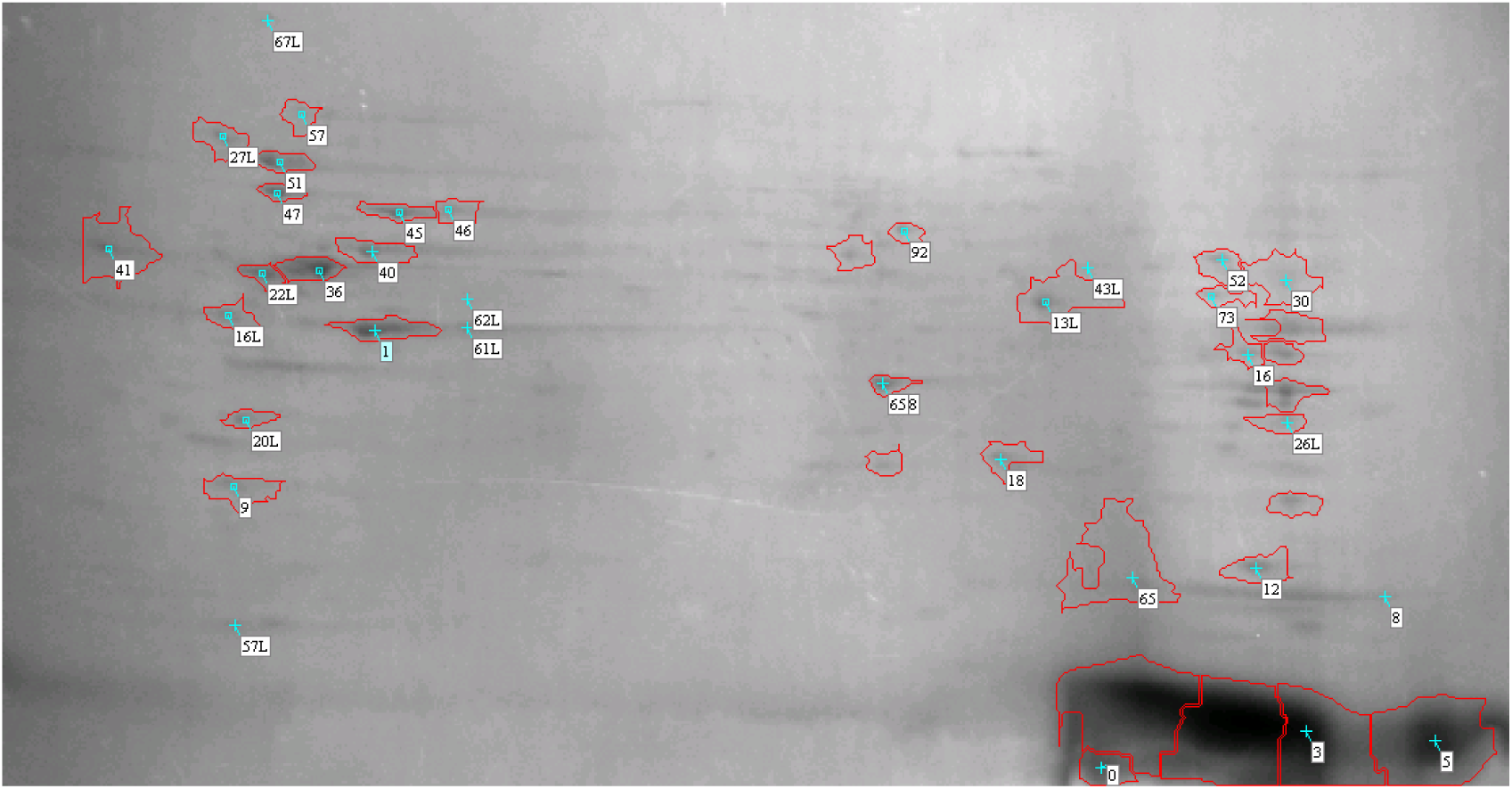
Protein profile of LH4+ Caps (4,0) treated with A2780^cisR^ cell line

**FIGURE 7.**
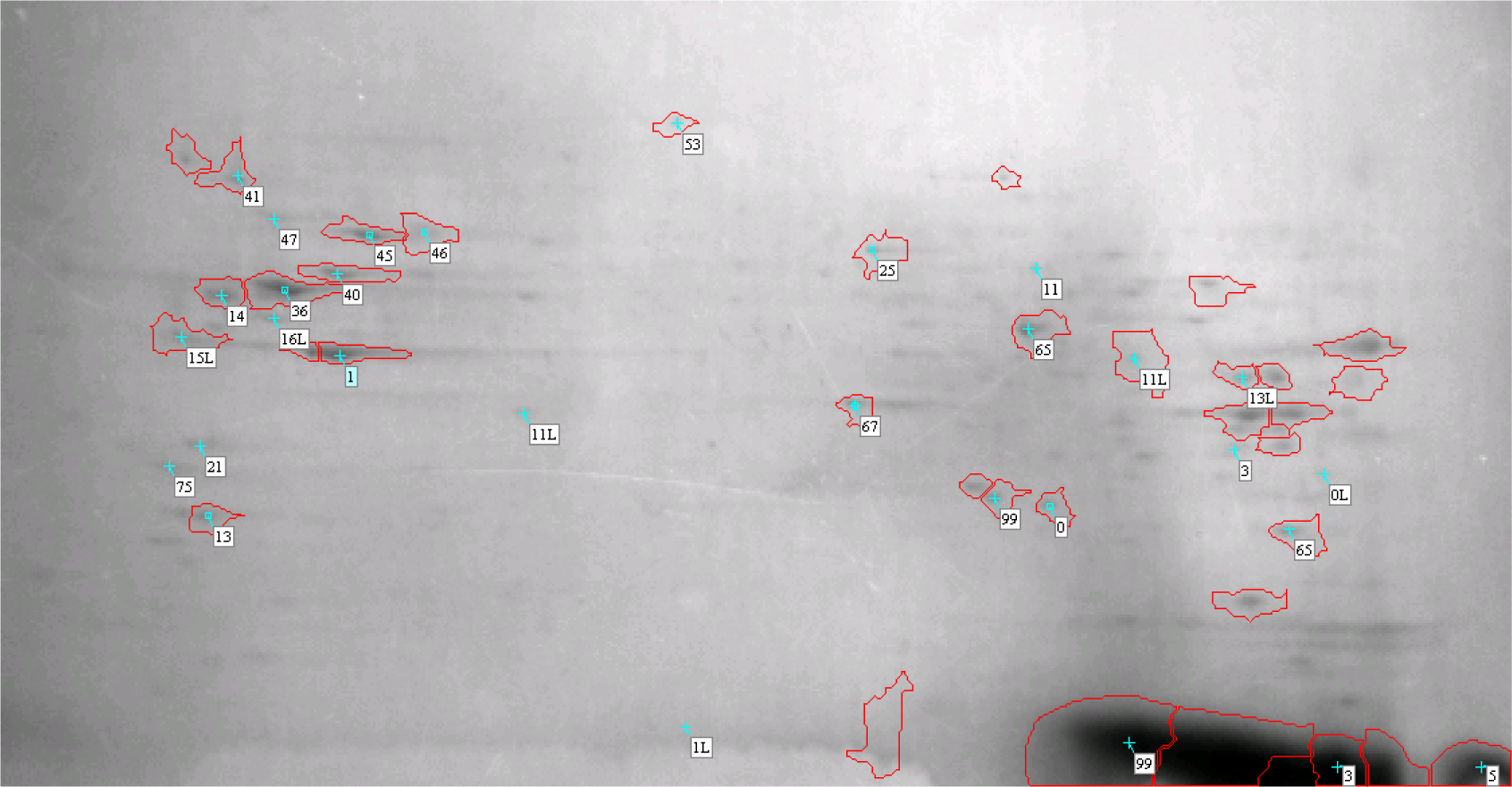
Protein profile of LH4+ Quer (0,0) treated with A2780^cisR^ cell line

A protein spot is considered to have undergone significant changes in expression if the fold change > 1.5 [18]. To determine the effect of drug treatment, expressions of selected 27 proteins in drug-treated A2780^cisR^ cell line were compared with the levels found in untreated A2780 and A2780^cisR^ cell lines, showing significant changes in expression (Figure 3 to 7 and Table 1). These 27 proteins with significant changes in expression were associated with cisplatin resistance in A2780^cisR^ cell line. Hence further analysis was done towards identifying these proteins (Table 2). To that end, these protein spots from the 2D-gel were excised and subjected to in-gel digestion with trypsin. Trypsin cleaved the proteins at C-terminal side of Lysine (K) and Arginine (R). Partial and fully fragmented proteins were scanned using MALDI-TOF/TOF MS analysis. Protein identification with details (Table 2), mass spectrum and matched peptides of 27 proteins were found from MALDI-TOF/TOF MS analysis (Supplementary Fig 1 to Fig 27).

**TABLE 1.**
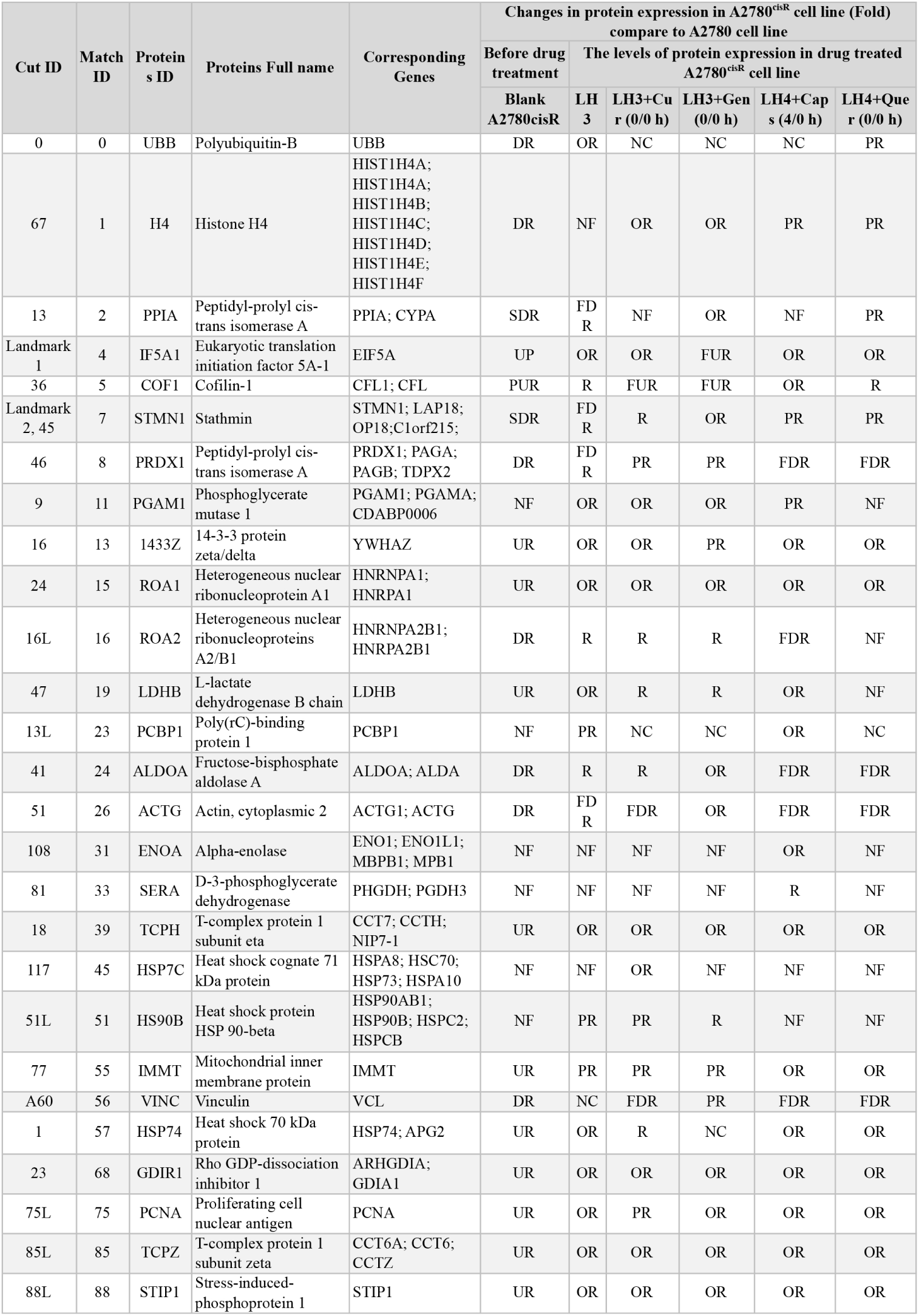
Changes in expression of selected protein spots in A2780^cisR^ cell line (down-or up-regulated) as compared to A2780 cell line before and after drug treatment. (DR= Down Regulated; UR=Up Regulated; SDR= Slightly Down Regulated; R= Restored; PR= Partially Restored; OR=Over Restored; FDR= Further Down Regulated; FUR=Further Up Regulated; NC= no change in expression; NF=not found, may be due to extreme down regulation. NB: A protein spot is considered to have undergone significant changes in expression if the fold change >1.5 [18]

**TABLE 2.**
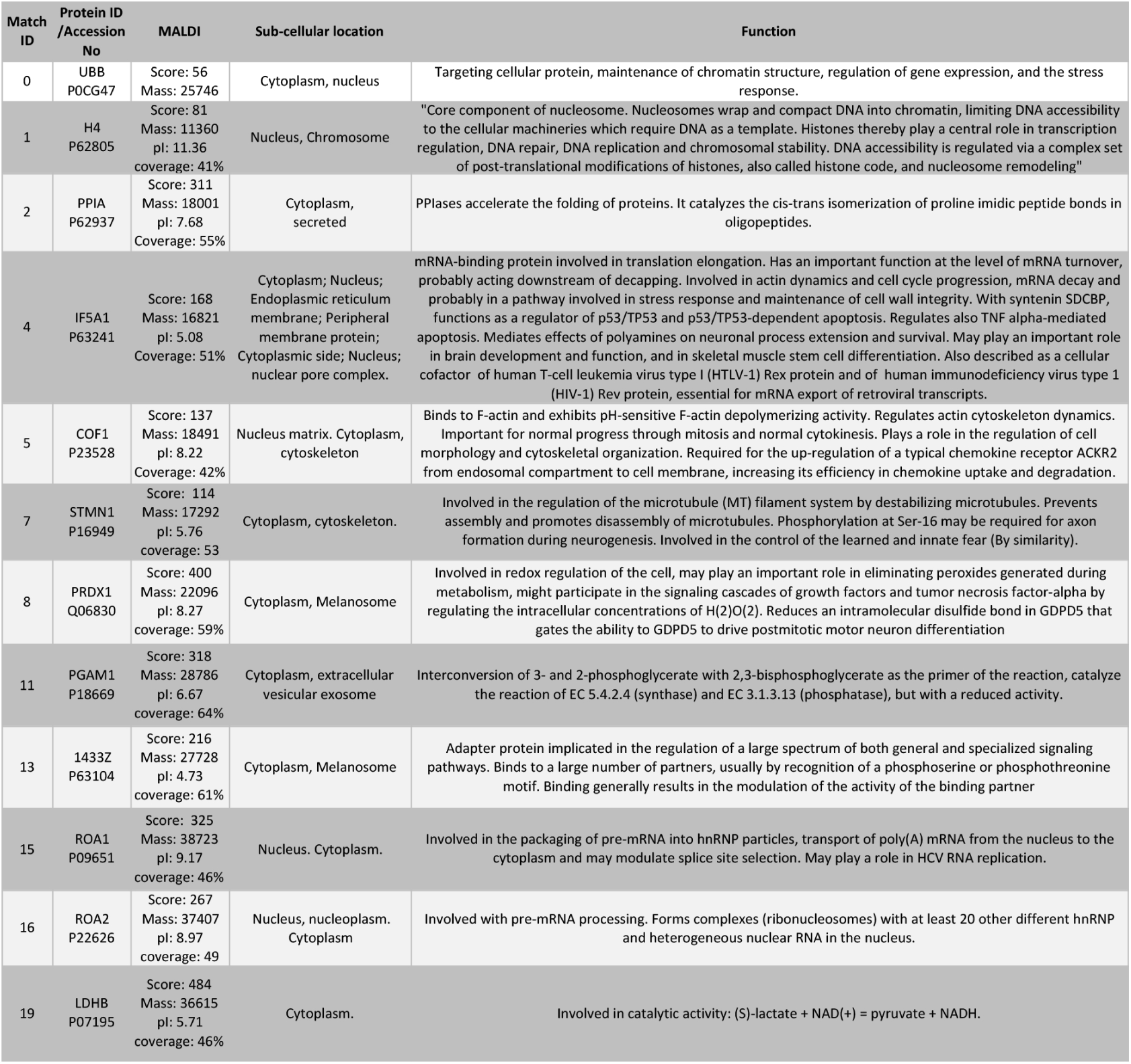
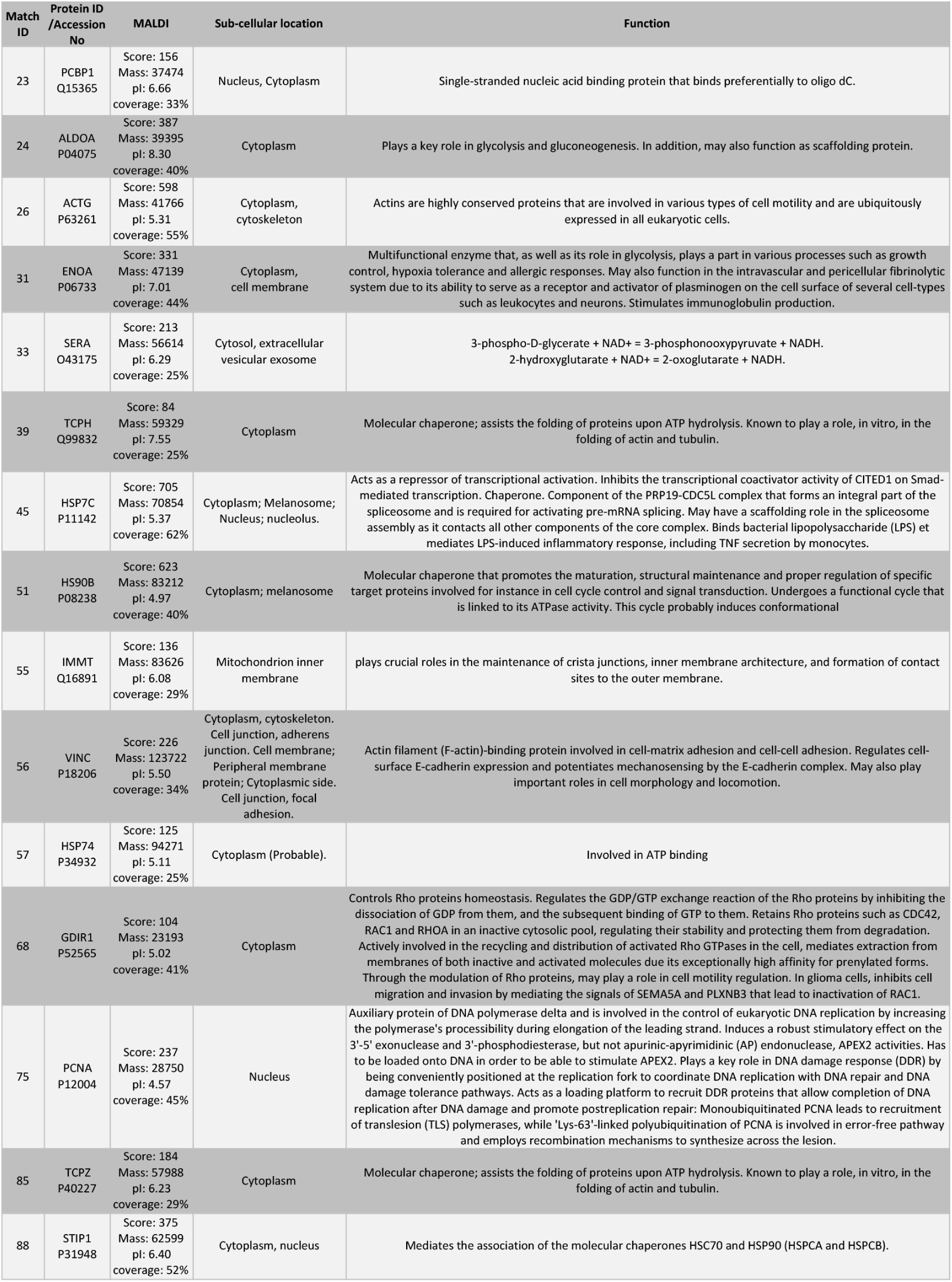
Changes in expression of selected protein spots in A2780^cisR^ cell line (down-or up-regulated) as compared to A2780 cell line before and after drug treatment. (DR= Down Regulated; UR=Up Regulated; SDR= Slightly Down Regulated; R= Restored; PR= Partially Restored; OR=Over Restored; FDR= Further Down Regulated; FUR=Further Up Regulated; NC= no change in expression; NF=not found, may be due to extreme down regulation. NB: A protein spot is considered to have undergone significant changes in expression if the fold change >1.5 [18]

6 clinical factors (including age) and RNASeq gene expression data for were employed in our analysis. All clinical factors were categorical except age. Age distribution of patients is shown in Figure 8. Average age of the patients at the time of their diagnoses was 59.7 years, with a range between 27 and 89 years old. The descriptive summary statistics of these factors shown in Table 3. From Table 2, we observe that the OC stage variable has the highest number (10) of categories and 71.03% of all patients are of stage IIIC. The next largest group is stage IV with 15.36% cases. There are 8 categories for histology type with the highest percentage of patients (84.67%) from was G3, i.e., poorly differentiated, while the second largest group (12.02%) of patients had moderately differentiated grade. Regarding ethnicity, most patients (85.27%) are from the European-caucasian descent population, while African American is next largest group with 5.89%. In the case of OC anatomical site, most (72.53%) women had bilateral cancers, while cancer patients with left and right ovary are 14.65% and 12.82% respectively. Most women had cancer in the ovary itself (99.13%), the remaining minor percentage of women had tumours located in omentum and/or peritoneum.

**TABLE 3.**
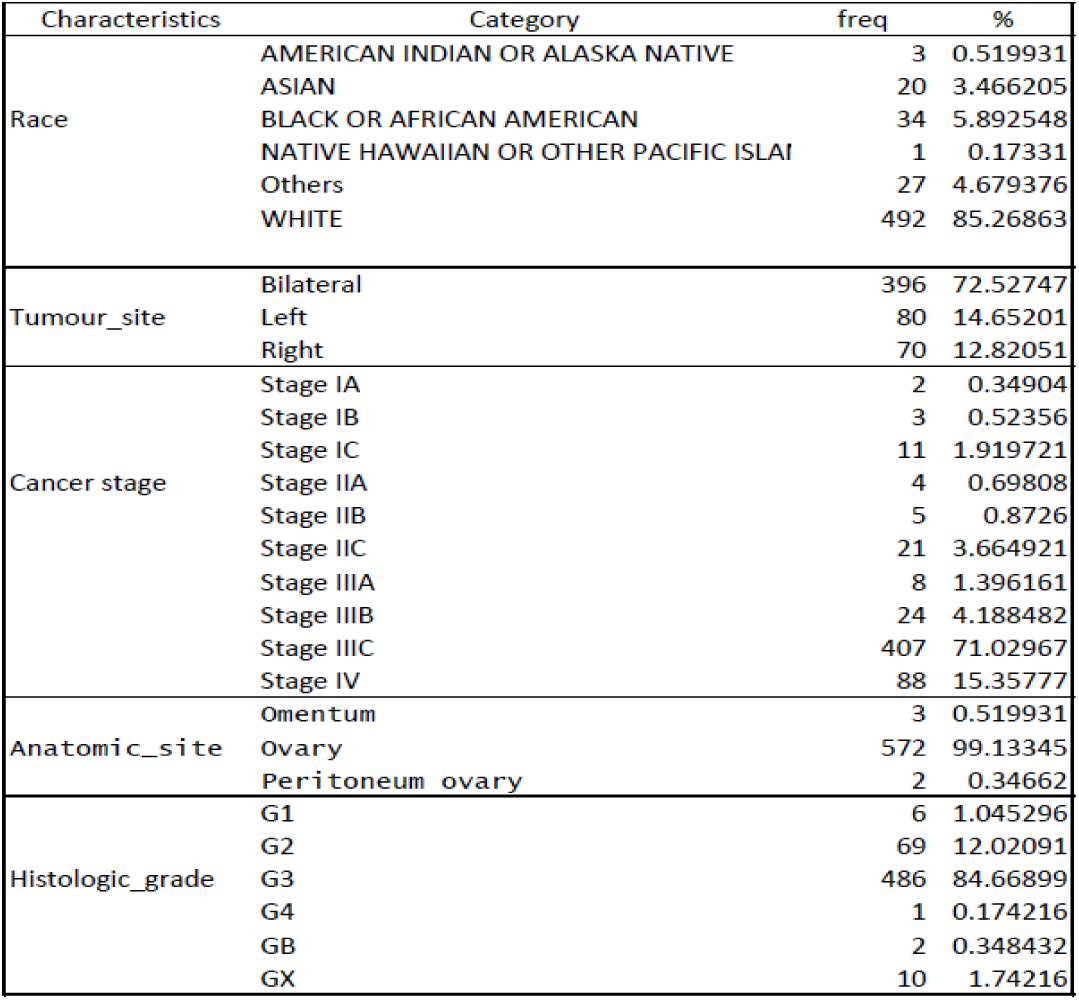
Descriptive Statistics of Clinical Predictors

| Characteristics | Category | freq | % |
| --- | --- | --- | --- |
| Race | AMERICAN INDIAN OR ALASKA NATIVE | 3 | 0.519931 |
|  | ASIAN | 20 | 3.466205 |
|  | BLACK OR AFRICAN AMERICAN | 34 | 5.892548 |
|  | NATIVE HAWAIIAN OR OTHER PACIFIC ISLAI | 1 | 0.17331 |
|  | Others | 27 | 4.679376 |
|  | WHITE | 492 | 85.26863 |
| Tumour_site | Bilateral | 396 | 72.52747 |
|  | Left | 80 | 14.65201 |
|  | Right | 70 | 12.82051 |
| Cancer stage | Stage IA | 2 | 0.34904 |
|  | Stage IB | 3 | 0.52356 |
|  | Stage IC | 11 | 1.919721 |
|  | Stage IIA | 4 | 0.69808 |
|  | Stage IIB | 5 | 0.8726 |
|  | Stage IIC | 21 | 3.664921 |
|  | Stage IIIA | 8 | 1.396161 |
|  | Stage IIIB | 24 | 4.188482 |
|  | Stage IIIC | 407 | 71.02967 |
|  | Stage IV | 88 | 15.35777 |
| Anatomic_site | Omentum | 3 | 0.519931 |
|  | Ovary | 572 | 99.13345 |
|  | Peritoneum ovary | 2 | 0.34662 |
| Histologic_grade | G1 | 6 | 1.045296 |
|  | G2 | 69 | 12.02091 |
|  | G3 | 486 | 84.66899 |
|  | G4 | 1 | 0.174216 |
|  | GB | 2 | 0.348432 |
|  | GX | 10 | 1.74216 |

**FIGURE 8.**
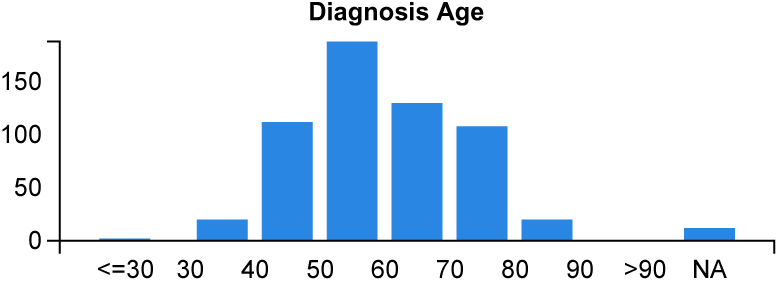
Distribution of Age at Onset of Diagnosis

### 3.1 Survival pattern for gene expression data

We have estimated survival function for altered and unaltered group for each of the 27 genes by applying product limit (PL) estimator. We then compared estimated survival function for altered and unaltered group using log rank test. Those genes for which there is statistically significant difference are shown below. The significant role for these genes is indicated by their p-values in differential survival pattern when comparing their expression level in two categories (altered and unaltered). Surprisingly p53 gene, which is known for its implication in cancer in general, has p-vlaue greater than 0.05. From Figure 9, we can see that patients having altered expression of ARHGDIA, CCT6A and HST1H4F genes are less likely to survive compared to the non-altered group. Note that, the red line in the graphs indicates normal gene expression and the blue line indicates the altered gene expression group.

**FIGURE 9.**
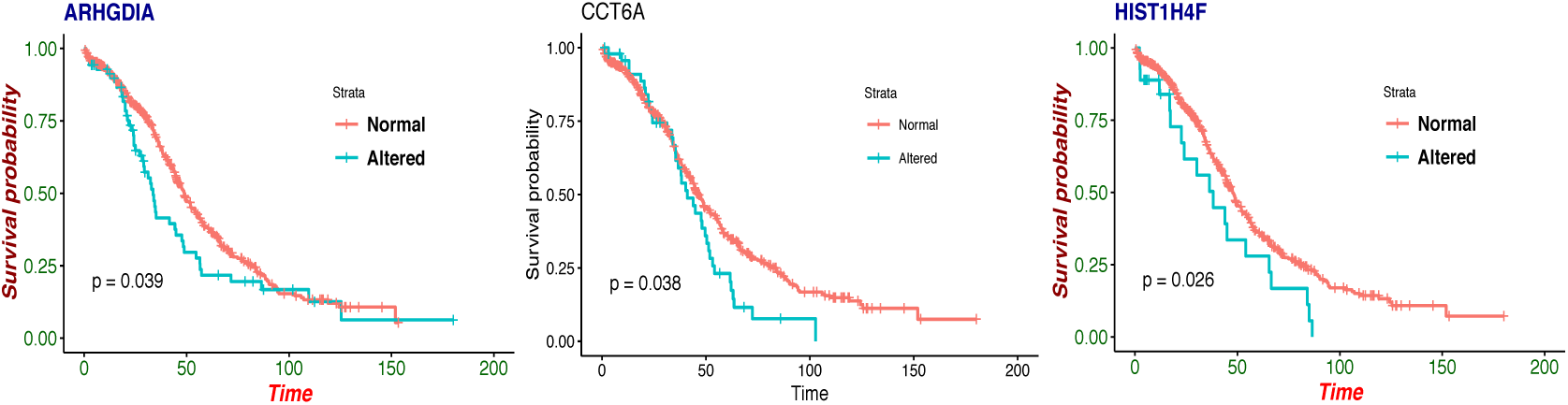
Survival pattern of altered and normal(non-altered) groups for ARHGDIA, CCT6A and HST1H4F

### 3.2 Modeling the hazard risk on the RNA-Seq data

One can measure the relative likelihood of risk of death in OC for each gene separately as well as for all genes simultaneously. Consequently one can determine which genes are most significant in case of survival of patients in OC. For these purposes, Cox PH regression model is used. We considered both univariate (separately for every gene) and multivariate (incorporating rest of the genes) for Cox PH model on each of the 27 genes. Table 3 shows the estimated Coefficients (*β*), with corresponding hazard ratios (HR), and p-values from those analyses.

We noted that the p-values for genes CD36 and KLK6 showed that their expression profile have statistically significant association with OC patient survival in both univariate and multivariate analyses. In contrast, genes SCGB2A and MEF2C showed significant association only in univariate analysis, while E2F1 showed significance only in multivariate analysis. Surprisingly, even though the KRAS gene has previously been shown by experimentation to have a significant role in OC ([32]), we did not find its significance in our analysis, both in univariate and multivariate analysis.

### 3.3 Pathway and functional correlation analysis of the significant genes

We observed that twenty significant pathways including microRNAs in cancer, bladder cancer, non-small cell lung cancer, and pancreatic cancer are associated with the significantly regulated genes for Ovarian Cancer. Genes associated with these pathways and corresponding p-values are presented in the table (see Table 4).

**TABLE 4.** Pathway associated with the selected 27 significantly associated genes with the Ovarian Cancer. Twenty two KEGG pathways were found using Enrichr for these genes. Genes associated with these pathways and corresponding Adjusted p-values are presented in this table.

| Kegg Pathways | Adjusted P-values | Genes |
| --- | --- | --- |
| Bacterial invasion of epithelial cells | 2.82E-05 | SEPT9;PIK3CA;CDH1;CTNNB1;VCL;ACTG1 |
| Viral carcinogenesis | 2.19E-04 | HIST1H4A;HIST1H4B;PIK3CA;HIST1H4C;HIST1H4D;YWHAZ;HIST1H4E;HIST1H4F |
| Adherens junction | 2.80E-04 | CDH1;ERBB2;CTNNB1;VCL;ACTG1 |
| Systemic lupus erythematosus | 7.09E-04 | HIST1H4A;HIST1H4B;HIST1H4C;HIST1H4D;HIST1H4E;HIST1H4F |
| Prostate cancer | 1.10E-03 | HSP90AB1;PIK3CA;ERBB2;AKT1;CTNNB1 |
| Pathogenic Escherichia coli infection | 9.80E-04 | CDH1;CTNNB1;YWHAZ;ACTG1 |
| HIF-1 signaling pathway | 1.26E-03 | PIK3CA;ERBB2;AKT1;ENO1;ALDOA |
| Central carbon metabolism in cancer | 1.83E-03 | PIK3CA;PGAM1;ERBB2;AKT1 |
| Glycolysis / Gluconeogenesis | 2.16E-03 | LDHB;PGAM1;ENO1;ALDOA |
| Alcoholism | 3.31E-03 | HIST1H4A;HIST1H4B;HIST1H4C;HIST1H4D;HIST1H4E;HIST1H4F |
| Proteoglycans in cancer | 5.65E-03 | PIK3CA;ERBB2;AKT1;RRAS2;CTNNB1;ACTG1 |
| Focal adhesion | 5.38E-03 | PIK3CA;ERBB2;AKT1;CTNNB1;VCL;ACTG1 |
| Fluid shear stress and atherosclerosis | 5.24E-03 | HSP90AB1;PIK3CA;AKT1;CTNNB1;ACTG1 |
| Regulation of actin cytoskeleton | 7.60E-03 | DIAPH2;PIK3CA;CFL1;RRAS2;VCL;ACTG1 |
| Leukocyte transendothelial migration | 1.26E-02 | PIK3CA;CTNNB1;VCL;ACTG1 |
| Thyroid hormone signaling pathway | 1.41E-02 | PIK3CA;AKT1;CTNNB1;ACTG1 |
| Mitophagy | 1.53E-02 | PRKN;UBB;RRAS2 |
| Rap1 signaling pathway | 2.52E-02 | PIK3CA;CDH1;AKT1;CTNNB1;ACTG1 |
| Estrogen signaling pathway | 2.44E-02 | HSPA8;HSP90AB1;PIK3CA;AKT1 |
| Antigen processing and presentation | 2.39E-02 | HSPA8;HSP90AB1;HSPA4 |
| Hepatitis C | 3.61E-02 | PIK3CA;AKT1;CTNNB1;YWHAZ |
| Fc gamma R-mediated phagocytosis | 3.66E-02 | PIK3CA;CFL1;AKT1 |
| Tight junction | 4.79E-02 | PCNA;HSPA4;ERBB2;ACTG1 |
| Progesterone-mediated oocyte maturation | 4.52E-02 | HSP90AB1;PIK3CA;AKT1 |
| Longevity regulating pathway | 4.86E-02 | HSPA8;PIK3CA;AKT1 |
| Hippo signaling pathway | 3.98E-02 | CDH1;CTNNB1;YWHAZ;ACTG1 |
| Hepatocellular carcinoma | 4.62E-02 | PIK3CA;AKT1;CTNNB1;ACTG1 |
| Hepatitis B | 4.21E-02 | PCNA;PIK3CA;AKT1;YWHAZ |
| Glycine, serine and threonine metabolism | 4.07E-02 | PGAM1;PHGDH |
| Glucagon signaling pathway | 4.98E-02 | LDHB;PGAM1;AKT1 |

We also performed the biological process ontology enrichment analysis (see table 5) of these identified significant genes using the Enrichr software tool. We found biological pathways that are associated with these significant genes as shown in Table 6.

**TABLE 5.** Gene Ontolgy terms with underlying Biological pathways associated with 27 significant genes in context of the Ovarian Cancer along. Eighteen Gene Ontology terms along with biological pathways, found significantly associated with these significant genes with adjusted p-values shown in this table.

| Go Terms | Pathways | Adjusted P-value | Genes |
| --- | --- | --- | --- |
| **Go Biological Process** |  |  |  |
| GO:0000731 | DNA synthesis involved in DNA repair | 1.46E-05 | RAD51D;PCNA;RAD51C;UBB;BRCA1;BRCA2 |
| GO:0071897 | DNA biosynthetic process | 1.62E-05 | RAD51D;RAD51C;BRCA1;BRCA2;PPIA |
| GO:0035635 | entry of bacterium into host cell | 1.35E-04 | UBB;CDH1;CTNNB1 |
| GO:0030260 | entry into host cell | 1.52E-04 | UBB;CDH1;CTNNB1;PPIA |
| GO:1904358 | positive regulation of telomere maintenance via telomere lengthening | 1.70E-04 | CCT6A;HNRNPA2B1;CCT7;HNRNPA1 |
| GO:0006986 | response to unfolded protein | 4.17E-04 | PRKN;HSPA8;HSP90AB1;HSPA4 |
| GO:0034620 | cellular response to unfolded protein | 7.91E-04 | PRKN;HSPA8;HSPA4 |
| GO:2000573 | positive regulation of DNA biosynthetic process | 1.05E-03 | CCT6A;HSP90AB1;CCT7;HNRNPA1 |
| GO:0061620 | glycolytic process through glucose-6-phosphate | 1.14E-03 | PGAM1;ENO1;ALDOA |
| GO:0061718 | glucose catabolic process to pyruvate | 1.14E-03 | PGAM1;ENO1;ALDOA |
| GO:0061621 | canonical glycolysis | 1.14E-03 | PGAM1;ENO1;ALDOA |
| GO:1902253 | regulation of intrinsic apoptotic signaling pathway by p53 class mediator | 1.30E-03 | PRKN;UBB |
| GO:0034329 | cell junction assembly | 1.36E-03 | CDH1;CTNNB1;VCL;ACTG1 |
| GO:0031396 | regulation of protein ubiquitination | 1.37E-03 | PRKN;HSP90AB1;UBB;AKT1;BRCA1 |
| GO:0035967 | cellular response to topologically incorrect protein | 1.42E-03 | PRKN;HSPA8;HSPA4 |
| GO:0034333 | adherens junction assembly | 1.73E-03 | CTNNB1;VCL |
| GO:1904869 | regulation of protein localization to Cajal body | 2.21E-03 | CCT6A;CCT7 |
| GO:0032212 | positive regulation of telomere maintenance via telomerase | 2.30E-03 | CCT6A;CCT7;HNRNPA1 |
| GO:1904851 | positive regulation of establishment of protein localization to telomere | 2.74E-03 | CCT6A;CCT7 |
| GO:1901214 | regulation of neuron death | 2.80E-03 | PRKN;UBB;AKT1;RRAS2 |
| **Go Cellular Component** |  |  |  |
| GO:1904813 | ficolin-1-rich granule lumen | 4.90E-04 | HSPA8;HSP90AB1;PGAM1;ALDOA;PPIA;VCL |
| GO:0060205 | cytoplasmic vesicle lumen | 6.29E-04 | HSPA8;HSP90AB1;PGAM1;ALDOA;PPIA;VCL |
| GO:0005657 | replication fork | 1.28E-03 | RAD51D;PCNA;RAD51C |
| GO:0000781 | chromosome, telomeric region | 3.34E-03 | HIST1H4A;RAD51D;PCNA;HNRNPA2B1;BRCA2 |
| GO:0101002 | ficolin-1-rich granule | 3.78E-03 | HSPA8;HSP90AB1;PGAM1;ALDOA;PPIA;VCL |
| GO:0005832 | chaperonin-containing T-complex | 5.43E-03 | CCT6A;CCT7 |
| GO:0005925 | focal adhesion | 8.16E-03 | HSPA8;CFL1;RRAS2;CTNNB1;PPIA;YWHAZ;VCL; |
| GO:0000784 | nuclear chromosome, telomeric region | 1.11E-02 | HIST1H4A;RAD51D;PCNA;BRCA2 |
| GO:0098687 | chromosomal region | 2.39E-02 | RAD51D;FMR1;HNRNPA2B1 |
| GO:0099513 | polymeric cytoskeletal fiber | 3.32E-02 | CCT6A;SEPT9;POF1B;CCT7;ACTG1 |
| GO:0034774 | secretory granule lumen | 4.30E-02 | HSPA8;HSP90AB1;PGAM1;ALDOA;PPIA;VCL |
| GO:0005694 | chromosome | 4.40E-02 | HIST1H4A;FMR1;BRCA1 |
| **Go Molecular Function** |  |  |  |
| GO:0045296 | cadherin binding | 4.67E-05 | SEPT9;HSPA8;HSP90AB1;CDH1;PRDX1;PCBP1;CTNNB1;ENO1;ALDOA;YWHAZ;VCL |
| GO:0003697 | single-stranded DNA binding | 7.07E-04 | RAD51D;RAD51C;PCBP1;BRCA2;HNRNPA1 |
| GO:0035198 | miRNA binding | 1.02E-03 | FMR1;HNRNPA2B1;HNRNPA1 |
| GO:0098505 | G-rich strand telomeric DNA binding | 3.34E-03 | HNRNPA2B1;HNRNPA1 |
| GO:0015631 | tubulin binding | 4.61E-03 | PRKN;RAD51D;FMR1;STMN1;BRCA1;BRCA2;ALDOA |
| GO:0043047 | single-stranded telomeric DNA binding | 4.68E-03 | HNRNPA2B1;HNRNPA1 |
| GO:0042623 | ATPase activity, coupled | 7.15E-03 | HSPA8;RAD51D;RAD51C;HSPA4 |
| GO:0070182 | DNA polymerase binding | 7.99E-03 | PCNA;HSP90AB1 |
| GO:0023026 | MHC class II protein complex binding | 7.99E-03 | HSPA8;HSP90AB1 |
| GO:0023023 | MHC protein complex binding | 1.10E-02 | HSPA8;HSP90AB1 |
| GO:0031625 | ubiquitin protein ligase binding | 2.73E-02 | PRKN;HSPA8;BRCA1;YWHAZ;VCL;ACTG1 |
| GO:0003723 | RNA binding | 2.90E-02 | EIF5A;HSPA8;HSP90AB1;FMR1;IMMT;ENO1;BRCA1;YWHAZ;CCT6A;STIP1;HIST1H4A;PRDX1;PCBP1;HNRNPA2B1;ALDOA;HNRNPA1;PPIA;SF1 |
| GO:0044389 | ubiquitin-like protein ligase binding | 3.30E-02 | PRKN;HSPA8;BRCA1;YWHAZ;VCL;ACTG1 |

Then we have investigated protein protein interaction network generated using STRING [38], a web-based visualization software resource as shown in figure 10. In this PPI network we have two groups of proteins in which fig 10a contains the previously identified ovarian cancer biomarkers and 10b contains our identified Oc drug resistance target. We have applied MCL cluster algorithm and found that our drug resistance target proteins are directly linked with the gold benchmark Oc biomarkers and exist in the same cluster. All of our genes are connected with each other through the PPI network as shown in 10.

**FIGURE 10.**
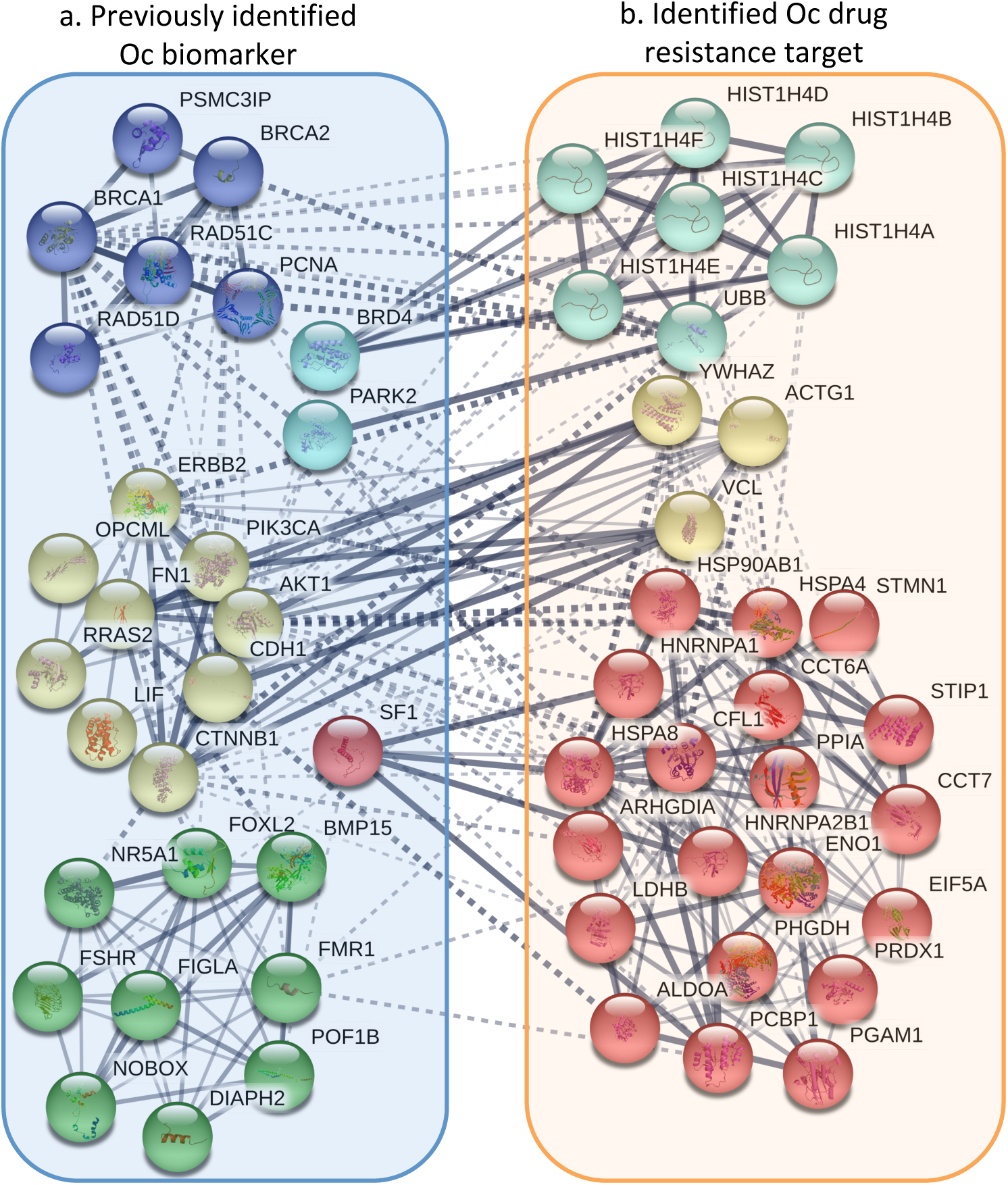
PPI network of 27 selected genes of OC

## 4 DISCUSSION

We used univariate Cox PH ratio analysis using RNASeq expression data, estimating the survival curve using product limit procedure and determining whether there is any statistically significant difference between the altered and un-altered groups using log rank test for each gene. This identified three significant genes (ARHGDIA, CCT6A and HST1H4F) in univariate analysis [30].

## 5 CONCLUSIONS

Proteomic studies have identified 122 proteins that were differentially expressed in A2780 and A2780^cisR^ cell lines. 27 of them were up-or down-regulated in A2780^cisR^ cell line due to drug trear data suggests that LH3 and LH4 are promising monofunctional platinums to overcome cisplatin resistance and differentially expressed proteins may be useful for further study of mechanisms of platinum resistance and screening of resistant biomarkers.

In this study, survival analysis of 577 OC patients revealed that out of 27 genes chosen for their previously idedntified involvment in OC, altered transcript levels of eight of these predicted reduced OC survival. Using product limit or Kaplan-Meier analyses, we found a significant difference in survival time between altered and non-altered genes. In cases of ARHGDIA, CCT6A and HST1H4F genes, patients with these altered expression of these three genes had significantly lower survival time than patients with non-alteration of these genes. It may be a candidate for therapeutic drug discovery itself or it may point the way to a crucial cell pathway that influences patient survival. Our approach can be used in case of other types of cancers to identify key genetic and clinical factors in patient survival.

## Abbreviations

OC: Ovarian Cancer
TCGA: The Broad Institute Cancer Genome Atlas

